# Dynamic vessel adaptation in synthetic arteriovenous networks

**DOI:** 10.1101/429027

**Authors:** Thierry Fredrich, Michael Welter, Heiko Rieger

## Abstract

Blood vessel networks of living organisms continuously adapt their structure under the influence of hemodynamic and metabolic stimuli. For a fixed vessel arrangement, blood flow characteristics still depend crucially on the morphology of each vessel. Vessel diameters adapt dynamically according to internal and external stimuli: endothelial wall shear stress, intravascular pressure, flow-dependent metabolic stimuli, and electrical stimuli conducted from distal to proximal segments along vascular walls. Pries et al. formulated a theoretical model involving these four local stimuli to simulate long-term changes of vessel diameters during structural adaption of microvascular networks. Here we apply this vessel adaptation algorithm to synthetic arteriovenous blood vessel networks generated by our simulation framework “Tumor-code”. We fixed the free model parameters by an optimization method combined with the requirement of homogeneous flow in the capillary bed. We find that the local blood volume, surface to volume ratio and branching ratio differs from networks with radii fulfilling Murray’s law exactly to networks with radii obtained by the adaptation algorithm although their relation is close to Murray’s law. In addition, we find that the application of the vessel adaptation algorithm to synthetic tumor vascularture does not lead to a stable radii distributions due to emerging short cuts between arteries and veins.

## 1 Introduction

Life depends crucially on the efficiency of transport networks and therefore it is essential to understand how they are functioning and why they are efficient [1]. In particular, the vasculature of all mammalians demonstrates how nature efficiently transports and distributes nutrients at minimal costs. Cecil Murray observed blood vessel networks and in 1926 and proposed a formula that described the observations on the thickness of the branches [2]. In her theoretical derivation of the formula she considered two energies with opposed dependency on the vessel radius r: the energy required to drive the flow and the energy required to maintain the metabolic demands of the surrounding tissue. To drive the flow, the system needs to overcome the viscous drag that is proportional to 1/*r*^4^ in case of Hagen-Poiseuille flow and to maintain the metabolic demands, a constant volume of “fresh” blood is required in a first approximation. By minimizing the sum of these two energies, she found that the sum of the cubes of the daughter vessel’s radii equals the cube of a parent vessel’s radius. In case of a simple junction with one mother vessel of radius *r*_*c*_ and two daughter vessels of radius *r*_*a*_ and *r*_*b*_, this reduces to:

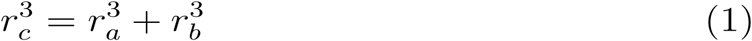

Since her assumptions are rather general, they hold true for the respiratory system of animals and insects, xylem in plants and diverse other kinds of transport networks as well [3]. Murray’s law is based on an optimality criterion and suggests the intriguing question how the vascularture of living organisms achieves this optimality? The blood vessel wall is formed by endothelial cells with a surrounding layer of smooth muscle cells. Because the muscular layer adjusts the vessel caliber continuously during growth, maturation and in response to exercise, wound healing and diseases, the vasculature is non-stationary with regulatory mechanisms that are still under investigation. While the vascular reactions to shear stress has been known for a long time [4, 5], recent studies showed that: 1) the shear stress is linked to the local intravascular pressure [6]; 2) the vascular diameter is controlled by the metabolic needs of the tissue (local oxygen partial pressure, metabolic signaling substances) [7, 8, 9, 10] and the metabolites remaining in blood after the exchange [11]; 3) electrical signals are conducted along the blood vessel wall by endothelial and smooth muscle cells via gap junctions [12, 13, 14] and 4) in case of hypertension, the vasculature switches to inward remodeling [15, 16].

Beginning in 1995 [6], Pries, Secomb, Gaehtgens et. al. published a series of papers [17, 18, 19, 16, 20] where they constructed a feedback loop between the discussed mechanisms and the blood vessel radii. Provided the resulting radius adaption dynamics leads to a stable state, this algorithm determines the blood vessel radii of a given network topology, in particular the relation between mother and daughter vessel radii at each junction without referring to Murray’s law.

In this manuscript we explore the application of the adaptation algorithm to in silico arteriovenous networks and analyze the emerging vessel radii and blood flow characteristics. Initially, the networks created by our in-house simulation framework Tumorcode [21] are consistent with Murray’s law. The vessel adaptation algorithm modifies each single vessel radius and we analyze the resulting distribution of radii. After validating our implementation by comparison with existing data in section 3.1, we optimize the free model parameters for a fixed vessel network in section 3.2 with respect to a maximally homogeneous blood flow in the capillary bed. In section 3.3, we compare the distribution of radii from the initial networks with the one emerging after the application of the vessel adaptation algorithm to them. Finally, the adaptation algorithm is applied to a simplified hypothetical network involving tumor growth induced modifications (section 3.5).

## 2 Methods

### 2.1 The adaptation algorithm

In [17], Pries et al. present a theoretical model to simulate vessel radii within arteriovenous networks. They constructed a feedback loop between the spatial position of the vessels (topology) and their biological functions including hydrodynamics, shear stress and metabolism of the surrounding tissue. For an elaborated review and some applications of the model see [22]. In the following, we describe the vessel adaptation algorithm of [17] that we use.

Considering a vessel of radius r during a time step Δ*t*, its adaption or change of a vessel radius Δ*r* = *S*_*tot*_ *r* Δ*t* is proportional to the sum of five signals 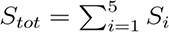 which are defined below.

#### S_1_ - Shear stress

Based on the correlation between wall shear stress and vessel radius, the adaptation algorithm reduces the vessel radius whenever the wall shear stress (*τ* _w_) falls below 1 dyne/cm^2^ = 10^*-*4^ kPa and extends the radius otherwise, i.e:

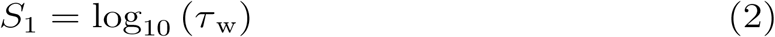

#### S_2_ - Blood pressure

In [6], the interaction of transmural pressure P and wall shear stress *τ*_*e*_ was studied, and a “pressure-shear” hypothesis was proposed

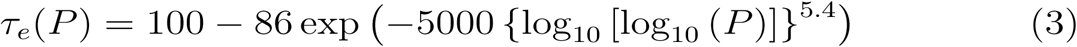

The functional form of the shear-pressure dependency is sigmoidally increasing from 14 dyne/cm^2^ for pressures of 10 mmHg to 100 dyne/cm^2^ for pressures at 90 mmHg (equation 3). With this

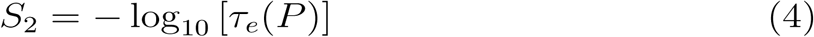

#### S_3_ - Metabolic demand

Similar to the idea of Murray, the metabolic demand needs to be maintained at all times. Since oxygen is transported as cargo by the red blood cells (RBC), a constant volume flow of RBCs is required to sustain a constant oxygen supply. Given the volume flow of blood Q with hematocrit H, the volume flow of RBCs is *HQ* and maintained at a reference value Q_ref_ by the metabolic signal S_3_.

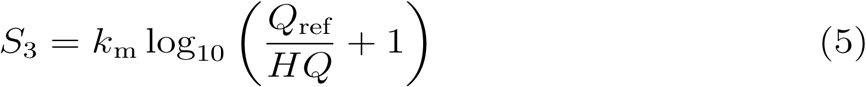

The definition of S_3_ results in a strictly positive signal where the amplitude increases stronger for vessels with a lower flow rate (*HQ* < *Q*_ref_) compared to vessels with a higher flow rate (*HQ* > *Q*_ref_) than the reference flow Q_ref_. In contrast to the biologically motivated reference flow, k_m_ is an unknown model parameter (compare table 1).

**Table 1:** Parameters of the adaptation model. The values in the right column are used in [17].

| signal | free | fixed |
| --- | --- | --- |
| $S_3$ | $k_m$ | $Q_{ref} = 40 \text{ nl/min}$ |
| $S_4$ | $k_c$ | $L = 1500 \text{ } \mu\text{m},$<br>$S_0 = 40$ |
| $S_5$ | $k_s$ | |

#### S_4_ - Topological position

*S*_4_ is sensitive to the position of the vessel within the entire network structure. The signal propagates from the place where the vessels interact most with the surrounding tissue: the capillary bed to the most upstream vessel in the arterial branch and the most down-stream vessel in the venous branch. While the downstream propagation of metabolic waste to the draining vein is reasonable, the upstream signaling is less obvious, but recent studies confirmed that vessels communicate via electrical signals within their endothelial layer [12, 13, 14]. In principle the algorithm works with arbitrary number of vessels per intersection. For simplicity, we limit the discussion to Y-shaped intersections where a vessel c has two downstream connections (to vessel a of length *x*_*a*_ and to vessel b of lenght *x*_*b*_) in an arterial branch and two feeding vessels (vessel a of length *x*_*a*_ and vessel b of lenght *x*_*b*_) in a venous branch. The conductive or topological signal is calculated in a recursive fashion by the weighted sum of the metabolic signals (S_3_) along the direction of information propagation (equation 6). The weighting factor is proportional to the exponential of the normalized length of the vessel segments 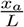 and 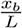.

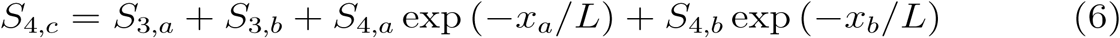

Since the recursive summation might result in excessive values, the signal is damped by a reference value *S*_0_ such that the complete conductive signal for a single vessel becomes:

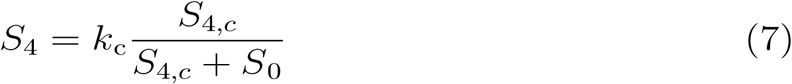

Note that the constant *k*_c_, the reference value *S*_0_ and the normalization length *L* are unknown model parameters (compare table 1).

#### S_5_ - Shrinking tendency

The introduction of a strictly positive metabolic (S_3_) and conductive signal (S_4_) necessitates a negative counterbalance. Its biological motivation is the tension in the endothelial layer which leads to a contraction of the vessel radius in the absence of all other signals. In the formula of the adaptation model this is embedded by subtracting a overall constant *k*_s_.

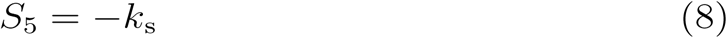

### 2.2 The artificial blood vessel networks

We construct artificial 3D arteriovenous networks by means of our simulation framework Tumorcode [21]. We report the complete methodology in [23] and summarize only the main steps here. First, at least one arterial and one venous root node have to be distributed in space and assigned with a hydrodynamic boundary condition (either a blood pressure value or a flow rate) before straight or Y-shape elements are appended until the arterial and venous branch merge by the capillaries. Second, the position of the capillaries within the volume is optimized towards a homogeneous capillary volume density using a hierarchical iteration scheme. For the scope of this paper, we consider only a root node configuration (RC) where one artery is opposite to one vein on the same axis (figure 2, compare also RC5 in figure 4 of [24]) and a single hierarchical iteration. Since the construction algorithm in the Tumorcode fixes the radii of all capillaries (2.5 *µ*m in presented case) and determines the radii of all intermittent vessels between root nodes and capillaries by Murray’s law, all radii at the intersections fulfill Murray’s law by construction. Therefore we label them **M-networks** in contrast to the **A-networks** which we obtain by applying the adaptation algorithm to the **M-networks**.

**Figure 1:**
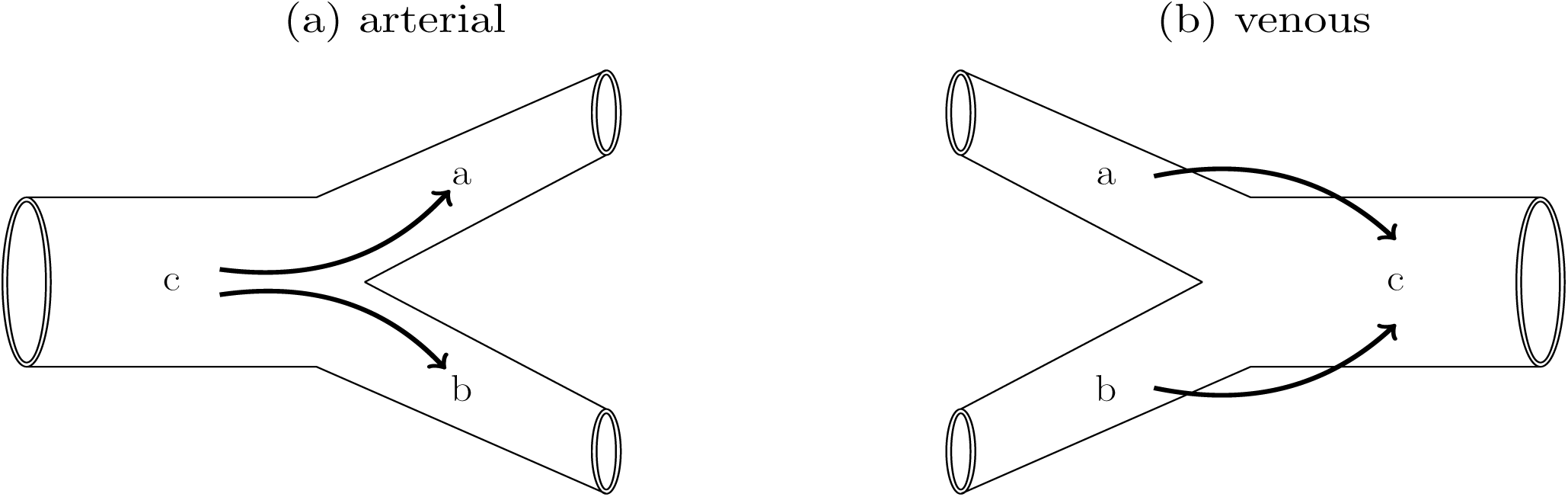
Recursive propagation of the conductive signal at a vessel junction. Left: arterial case. Right venous case. Arrows indicate the direction of information propagation. See equation 6.

**Figure 2:**
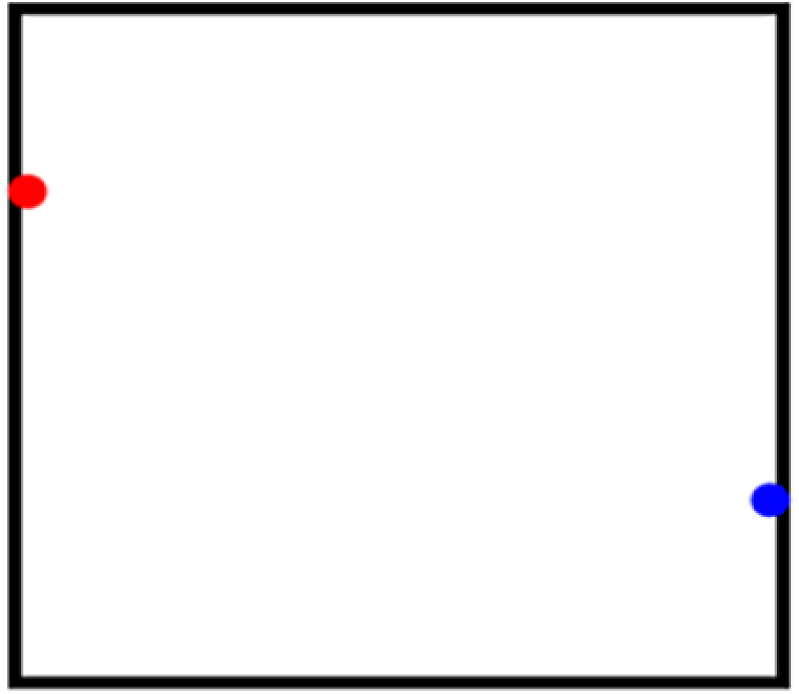
Schematic root node configuration (RC) for construction the arteriovenous networks. Red: arterial root, blue: venous root.

**Figure 3:**
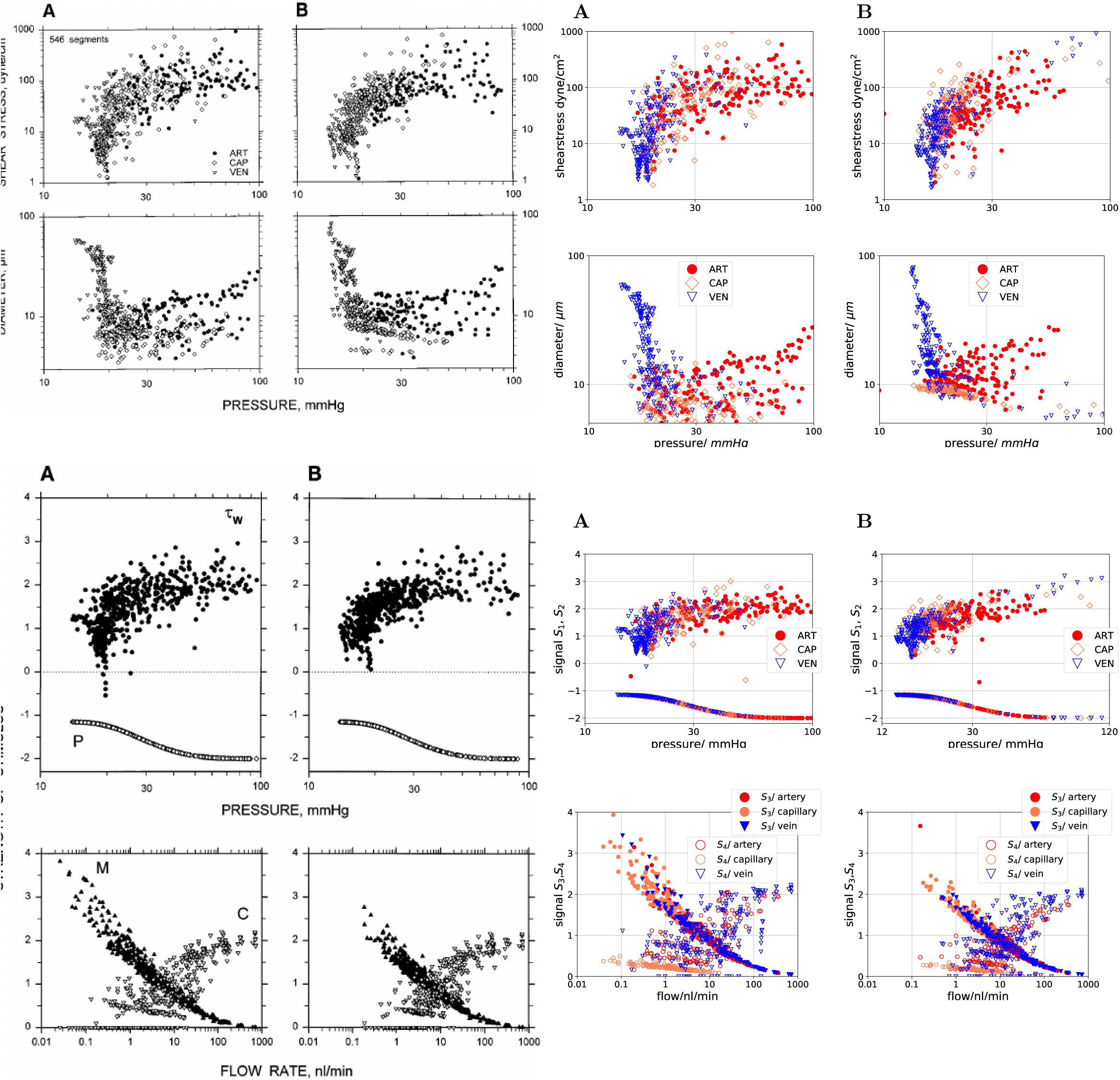
Left column (black and white): reprint of Figure 4 and 5 from [17] with permission. Right column (colored): reproduced by our implementation/ software. (A): values obtained with experimentally determined vessel diameters, (B): values obtained by simulated vessel adaptation. Top and second from top row: Distribution of shear stress and diameter for a rat mesentery network with 546 segments (162 arterioles, 167 capillaries and 217 venules). Hemodynamic parameters were calculated using the network flow model based on measured network morphology and topology. Last before bottom row: hydrodynamic stimuli derived from shear stress (*τ*_*w*_) and blood pressure (P) are plotted as functions10of pressure. Last row: the metabolic and conducted stimuli (C) are plotted as functions of flow rate, to show functional dependence of these stimuli.

**Figure 4:**
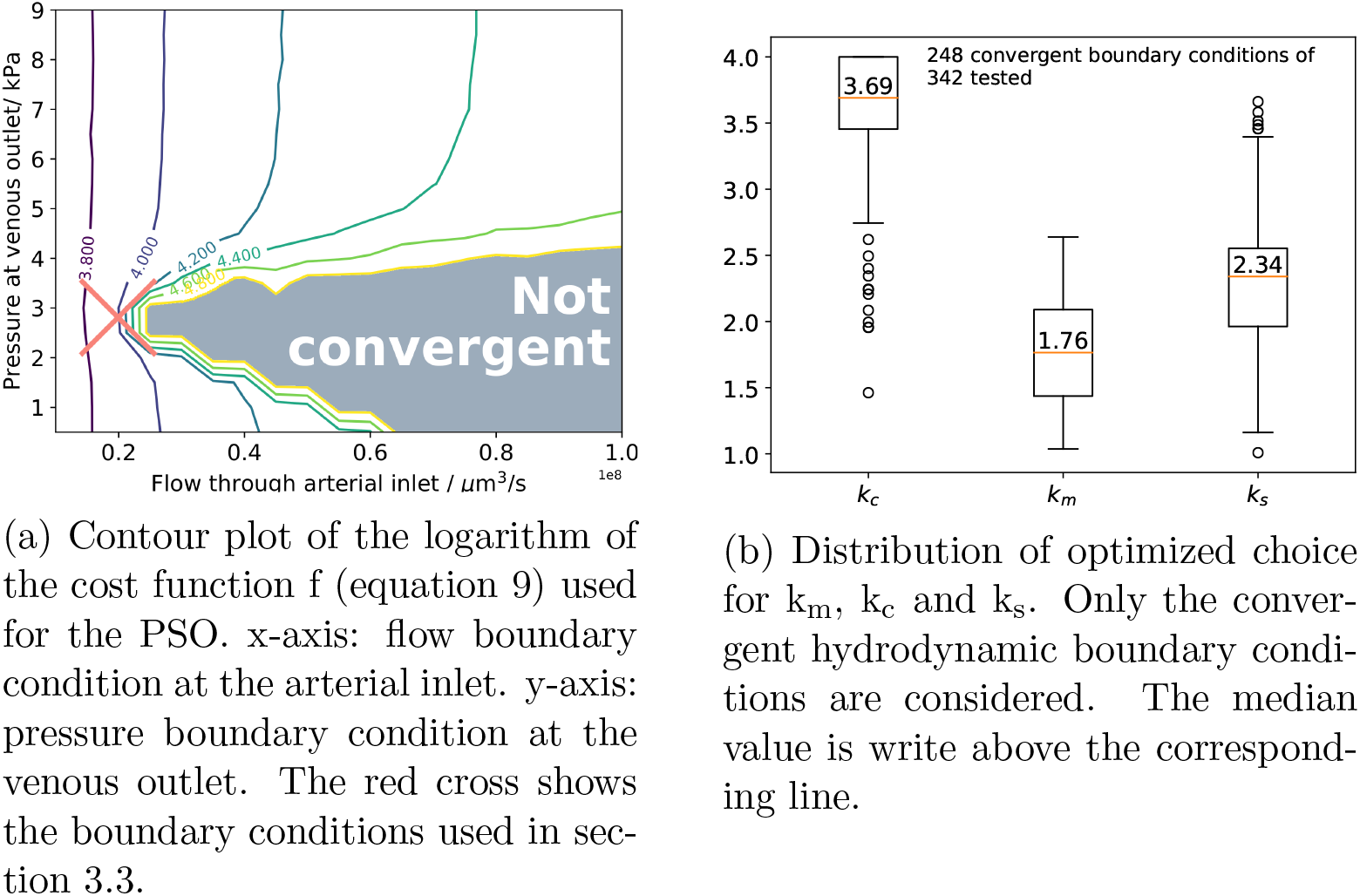
Results of the particle swarm optimization (PSO).

### 2.3 Parameter optimization

Inspecting section 2.1, we count six unknown model parameters as summarized in table 1. Considering the additional hydrodynamic boundary condition at the two root nodes, we end up with 8 degrees of freedom for the model. To find the unknown parameters, we presume that a properly working vasculature requires a homogeneous flow inside the capillary bed to maintain the nutrient exchange and apply a computational optimization technique where the variance of the volume blood flow inside all capillaries (*Q*_*capillaries*_) is used as cost function

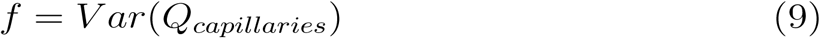

We used particle swarm optimization (PSO) [25] which minimizes the cost function f defined in equation 9. Although there is no guarantee to find the best solution, PSO finds at least a “robust” solution i.e. a solution that is robust against varying the initial conditions. In our case, the initial conditions are the values of the parameters k_m_, k_c_ and k_s_. During the PSO, we vary these parameters in the range of 0.5 to 4.0 for each fixed set of hydrodynamic boundary conditions. The hydrodynamic boundary conditions are the arterial inlet flow and the venous outlet pressure which we also varied in a systematic way (see supplemental text).

The values of *Q*_ref_ = 40*nl/min, L* = 1500*µm* and *S*_0_ = 20 were kept fixed at the values reported in [17].

The exact numbers, computational details and the implementation is reported in a supplemental text to this manuscript.

### 2.4 Definition of Biophysical Quantities

Given a blood vessel network inside a volume V that consists of N tubes with radii *r*_*i*_ and lenghts *l*_*i*_, two quantities of interest are the regional blood volume (rBV) and the surface to volume ratio (s2v).

#### 2.4.1 Regional Blood Volume (rBV)

The regional blood volume measures the percentage of blood within the volume.

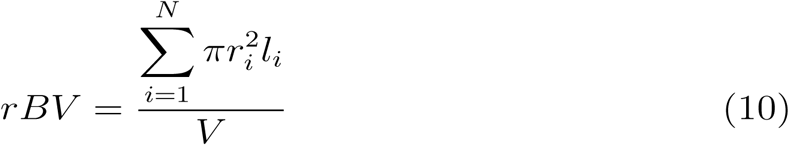

#### 2.4.2 Surface to volume ratio (s2v)

The surface to volume ratio measures the ratio between the surface of an object and the volume included by this surface. Since nutrient exchange happens across the vessel surface, the exchange is proportional to the surface and therefore the s2v is an interesting quantity.

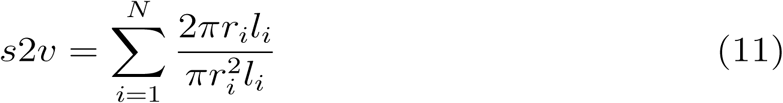

## 3 Results

### 3.1 Reproducing available data

We applied our implementation of the adaptation algorithm to a experimentally determined rat mesentery network. Since the hydrodynamic boundary conditions were measured, we incorporated them accordingly. The same procedure was done in [17] (Network I). Unfortunately, the used time step Δ*t* is not documented. We used Δ*t* = 0.1 and the other adaptation parameters in table 2. To verify our implementation, we reproduced two figures comparing the shear stress, vessel diameters, and the adaptation signals S_1_, S_2_, S_3_ and S_4_ before and after application of the algorithm.

**Table 2:**
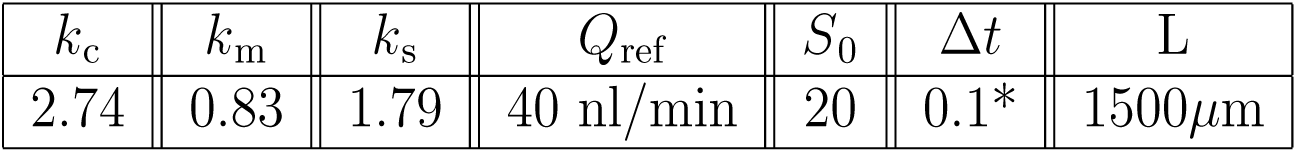
Parameters used to adapt vessel radii of rat mesentery network with Tumorcode. Parameter labeled by * is not shown in [17].

| $k_c$ | $k_m$ | $k_s$ | $Q_{\text{ref}}$ | $S_0$ | $\Delta t$ | L |
| --- | --- | --- | --- | --- | --- | --- |
| 2.74 | 0.83 | 1.79 | 40 nl/min | 20 | 0.1* | 1500$\mu\text{m}$ |

Due to the lack of quantitative data in [17] we cannot compare directly, but our figures are in qualitative agreement with the original (figure 3). For the hydrodynamics (top row and second from top row of figure 3), we observe that the shear stress saturates with increasing blood pressure (top row) and the diameter varies for veins in a broader range than for the arteries with the same pressure (second row). The adaptation tends to increase the radius of the capillaries (coral labeled points in rightmost image of second row). For the distribution of the adaptation signals S_1_ and S_2_ (last before bottom row of figure 3), we find vessels in the adapted network with a higher blood pressure than 100 mmHg and therefore we extended the pressure axis (x-axis) to 120 mmHg. For the distribution of the adaptation signals S_3_ and S_4_ (bottom row of figure 3), we find that adaptation shifts the spectrum towards increased flow. In agreement with model assumptions, the metabolic signal/ demand (S_3_) is higher for low flow segments and the conductive signal (S_4_) increases for segments with higher flow that are more up or downstream respectively.

### 3.2 Single topology, varying boundary conditions

As already discussed in [17], there are two possible outcomes when applying the adaptation scheme in a repeated manner: either one reaches an steady state where the changes of radii become arbitrary small with each iteration or the radii changes increase to unrealistic values which also implies unrealistic values for pressure and flow, and is not further consider.

By the optimization procedure described in section 2.3, we identified the hydrodynamic boundary conditions and parameters that result in convergence of the adaption algorithm with a fixed network topology (see figure 4a) and achieved an optimal homogeneity in the blood flow of the capillaries. Interestingly, the distribution of optimization parameters (figure 4b) indicates the same relation between the parameters (k_m_ < k_s_ < k_c_) as found by Secomb et. al. in their three samples of rat mesentery networks [17].

To estimate the biological relevance of the optimized networks, we analyze two biophysical variables (section 2.4): the regional blood volume (figure 5a) and the surface to volume ratio (figure 5b). The total rBV varies within a physiological regime from 2% up to 10% (left column in figure 5a). To estimate the contributions from arteries, capillaries and veins to the total rBV, we segmented them in figure 5a.

**Figure 5:**
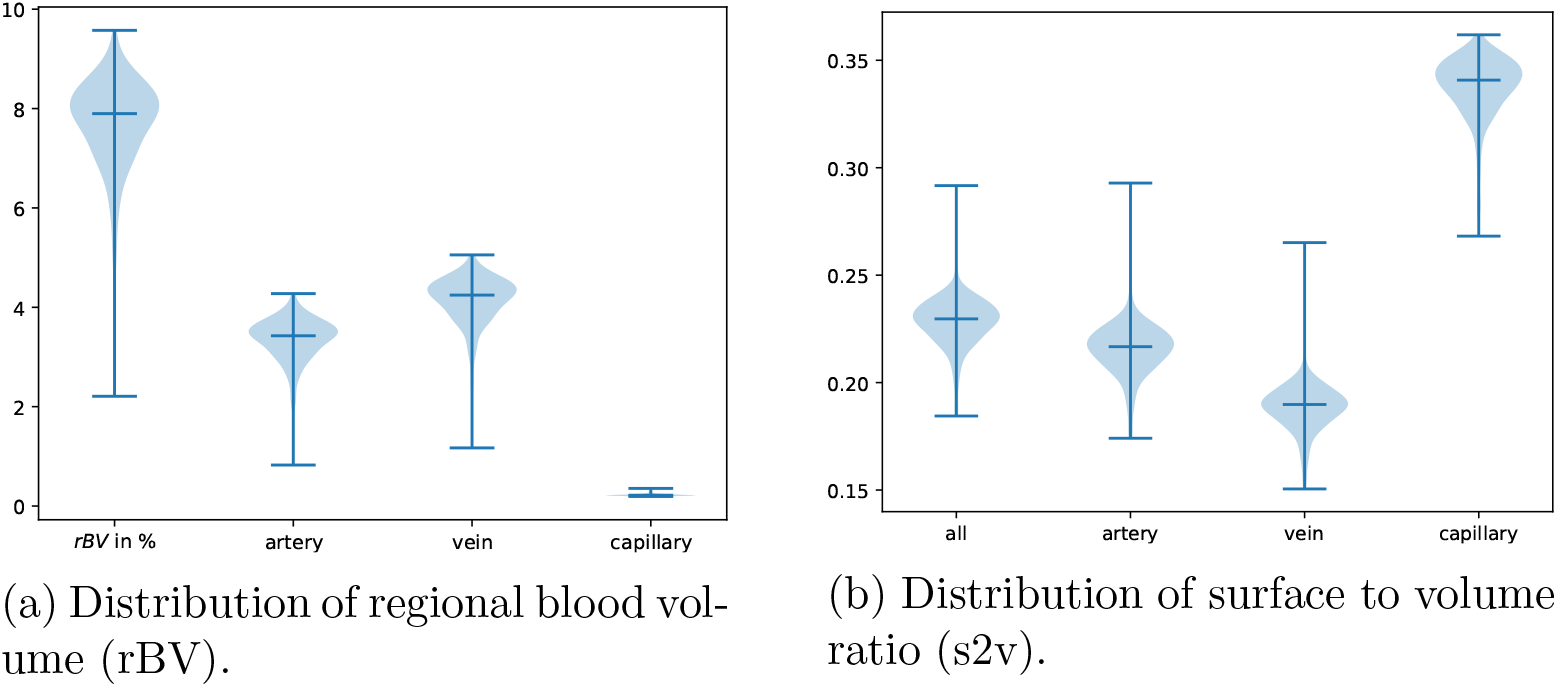
Biophysical variables influenced by the adaptation algorithm. Distribution is obtained from a single configuration of vessels in space (section 3.2), but different radii distributions obtained during particle swarm optimization. We considered the 248 convergent boundary conditions here.

The results for the surface to volume ratio are shown in similar fashion (figure 5b). As expected, the surface to volume ratio is highest for small capillaries, smaller for arteries and the smallest for the large veins.

### 3.3 Varying topology, fixed boundary conditions

In contrast to the previous section, we now 1) keep the hydrodynamic boundary conditions and adaptation parameters fixed and 2) vary the topological arrangement of the vessels. Based on the results of the PSO (red cross in figure 4a), we decided to use a pressure of 2.8 kPa at venous outlet and a volume flow rate of 2 ·10^7^ *µ*m^3^/s at the arterial inlet to create 10 independent realizations of arteriovenous networks (**M-networks**) of lateral size 1500 *µ*m (see section 2.2).

#### 3.3.1 Biophysical Quantities

We apply the adaptation algorithm with the parameter set listed in table 2 to the 10 samples. Hence we constructed 10 **A-networks** which we compare regarding their regional blood volume rBV (Figure 6a) and their surface to volume ratio s2v (Figure 6b) to the **M-networks**.

**Figure 6:**
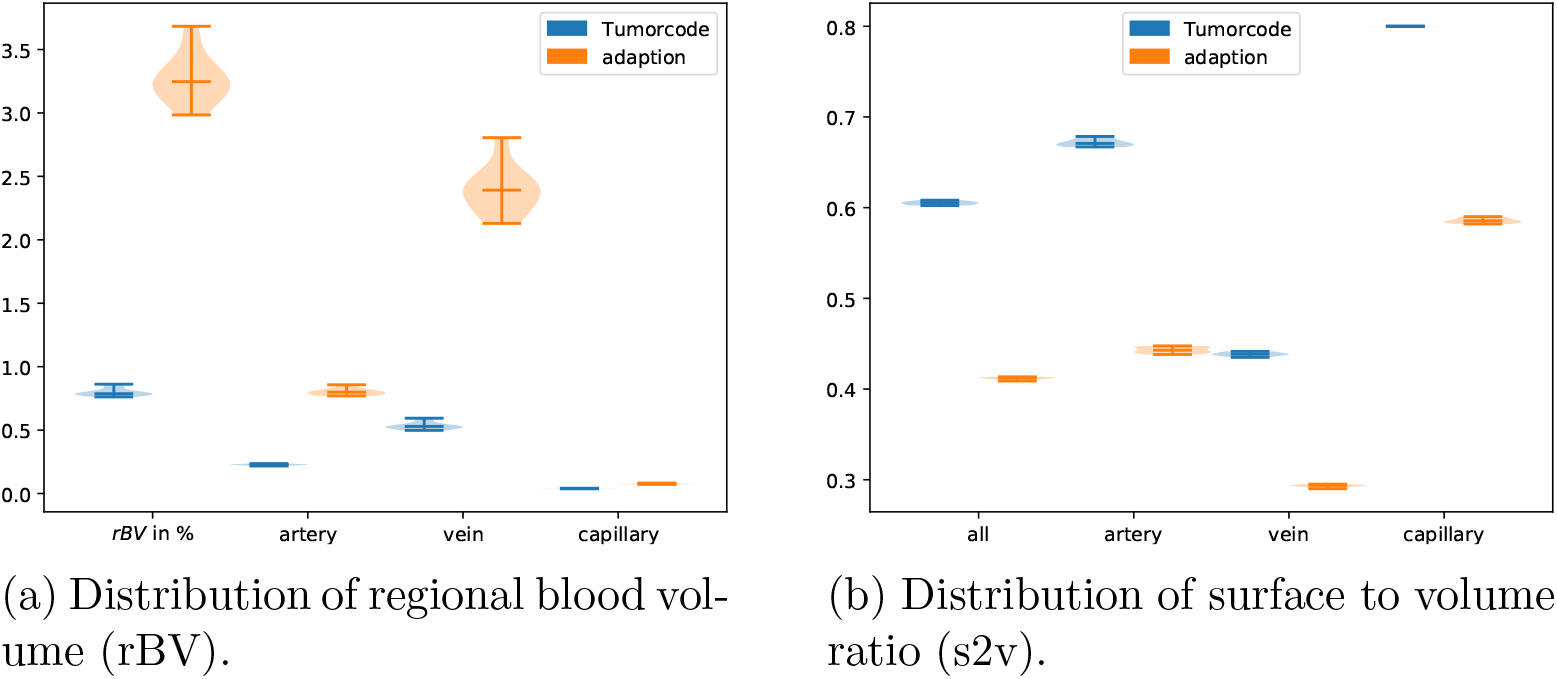
Biophysical variables influenced by the adaptation algorithm. Distribution is taken from 10 different arrangements of vessels in space with radii obtained by 1) construction by Tumorcode (blue) and 2) application of the adaptation algorithm with a fixed set of parameters (table 2, orange).

For the **A-networks**, the rBV is more than 3 time higher than for the **M-networks** (left column of figure 6a). While the contributions from arteries and veins to the overall rBV varies, the contribution from the capillaries are about the same (right column of 6a). Considering the surface to volume ratio (figure 6b), Tumorcode creates networks with a higher ratio resulting in a better biological efficiency. Note that the s2v ratio is highest for the capillaries in the M-networks.

We conclude that the optimization which arranges the capillaries in space during network construction in Tumorcode satisfies its purpose: namely to create a optimal environment for nutrient exchange.

#### 3.3.2 Adaptation signals

Similar to figure 3, we plot: a) the hydrodynamic characteristics (top and next to top row in figure 7), b) the hydrodynamic signal (last before bottom row in figure 7), and c) the conductive and metabolic signals (bottom row in figure 7) for one representative of the **A-networks** in figure 7. For the other nine networks, the plots look qualitatively similar.

**Figure 7:**
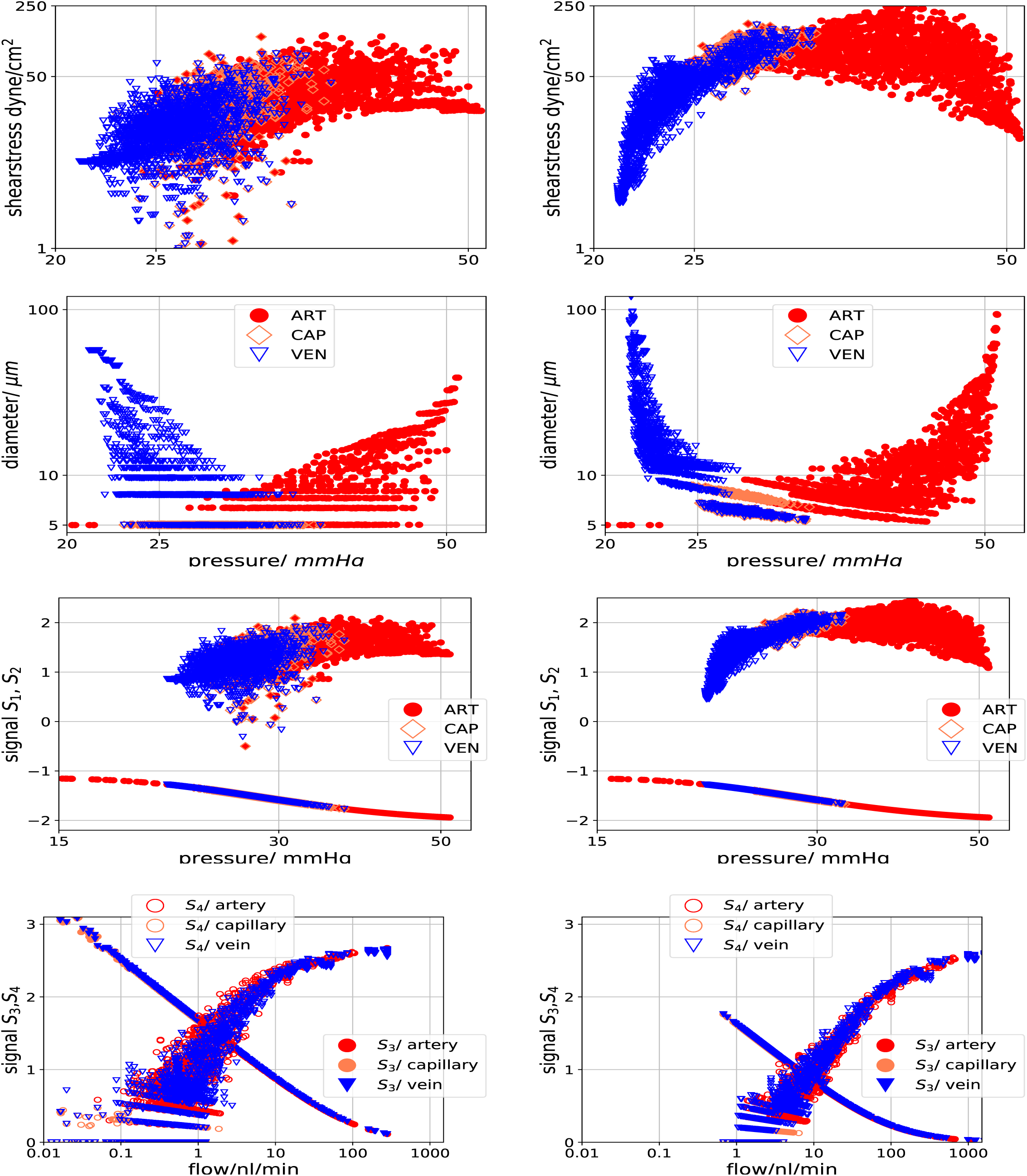
The characteristics for a single arteriovenous network. Left column: without adaptation. Right column: convergent adaptation with parameters listed in table 2. A graphical representation of this particular network is shown in figure 8. From top to bottom we compare the shear stress, the vessel diameter, the hydrodynamic signals (S1 and S2 of section 2.1) and the metabolic and conductive signal (S3 and S4 of section 2.1)

Before adaptation, there is no functional dependence of the shear stress on the pressure, and since the the signal S_1_ (positive values of last before bottom row) is calculated by the logarithm of the shear stress, there is also no functional dependence of S_1_ on the pressure. Although it is no clear relation, the dependence of the shear stress and the signal S_1_ on the pressure becomes more pronounced after the adaptation (right column of figure 7): veins approach a plateau with increasing pressure and the arteries with decreasing pressure. For the shear stress, the plateau is between 50-100 dyne/cm^2^ and for the signal S_1_ around 2.

The diameter depends on the pressure as expected for our networks: starting with the vessels of the smallest diameter (the capillaries), the diameter increases as we raise the pressure along the arterial branch and decrease the pressure along the venous branch (second row of figure 7). The negative values in the last before bottom row of figure 7 present the signal S_2_ (equation 4). Here we notice that the values for S_2_ are more restricted for veins (between −1.2 and −1.7) than for arteries (between −1.1 and −2.0). The metabolic and conductive signal (bottom row of figure 7) shift their maximum / minimum from 0.01 nl/min to increased flows at about 1 nl/min.

#### 3.3.3 Visual inspection

Although the distribution of radii changes from the **M-networks** (bottom left of figure 8) to **A-networks** (bottom right column of figure 8), the distribution of the blood pressure does no change because of the fixed boundary conditions (colors of top row in figure 8). Since the adaptation changes the radii, the flow needs to be modified accordingly! The shear force is proportional to the flow and therefore the distribution of the shear force looks similar to the distribution of flow as show in the center row of figure 8. In the bottom row of figure 8, we colored the vessel by their radii to highlight changes of the vessel radii by the adaptation algorithm.

**Figure 8:**
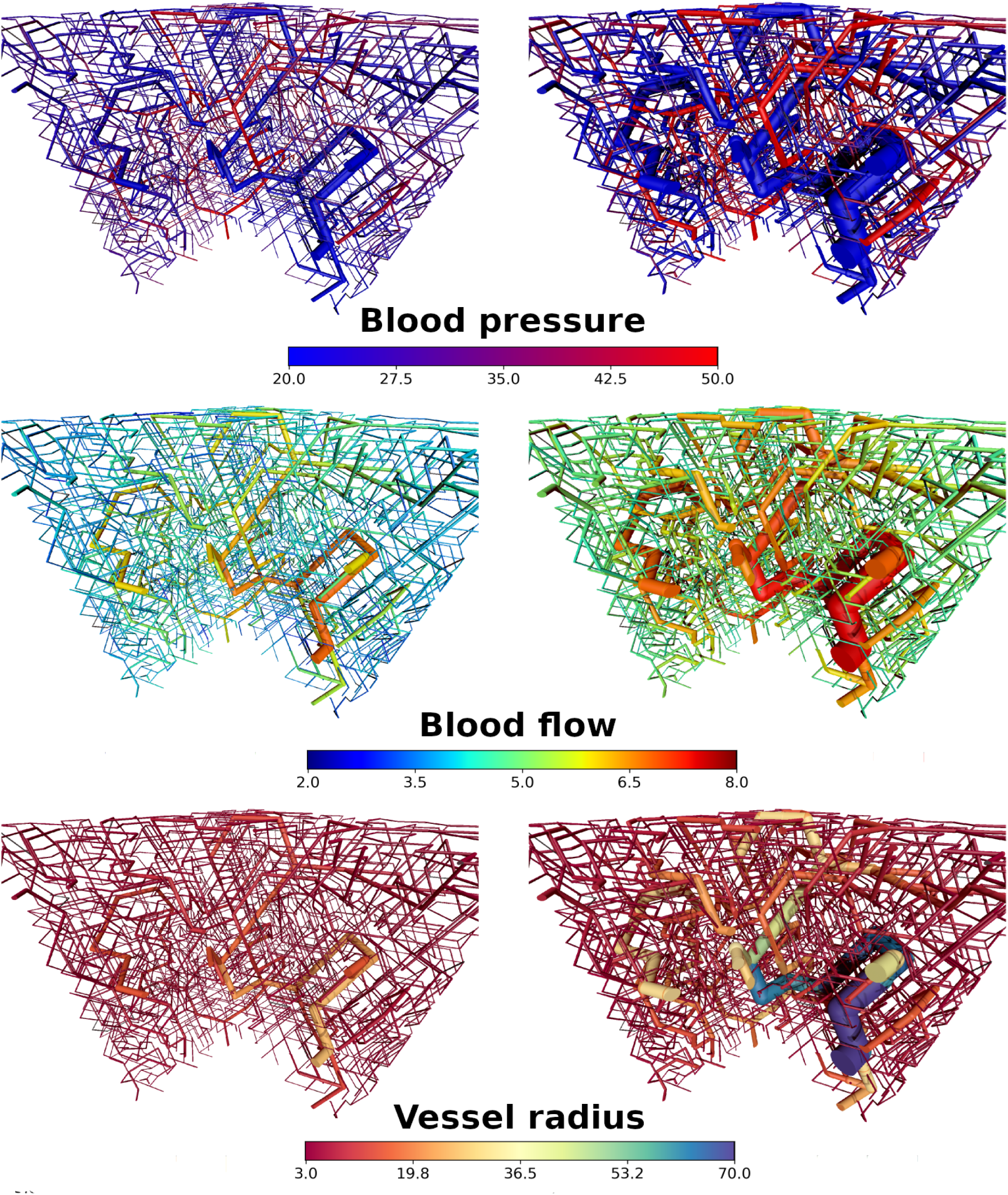
Color coded vessel properties for one representative of the ensemble of synthetic arteriovenous networks. Left column: radii distribution from Tumorcode (Murray’s law). Right column: radii distribution after adaptation. From top to bottom: pressure in mmHg, volume flow (logarithmic scale) in *µ*m^3^/s, radius in *µ*m.

Finally, figure 9 shows data that is only available in the adaption case (metabolic signal and conductive signal). In agreement with model assumptions, the metabolic signal is highest at the thin capillaries and the conductive signal is highest at the most distal points.

**Figure 9:**
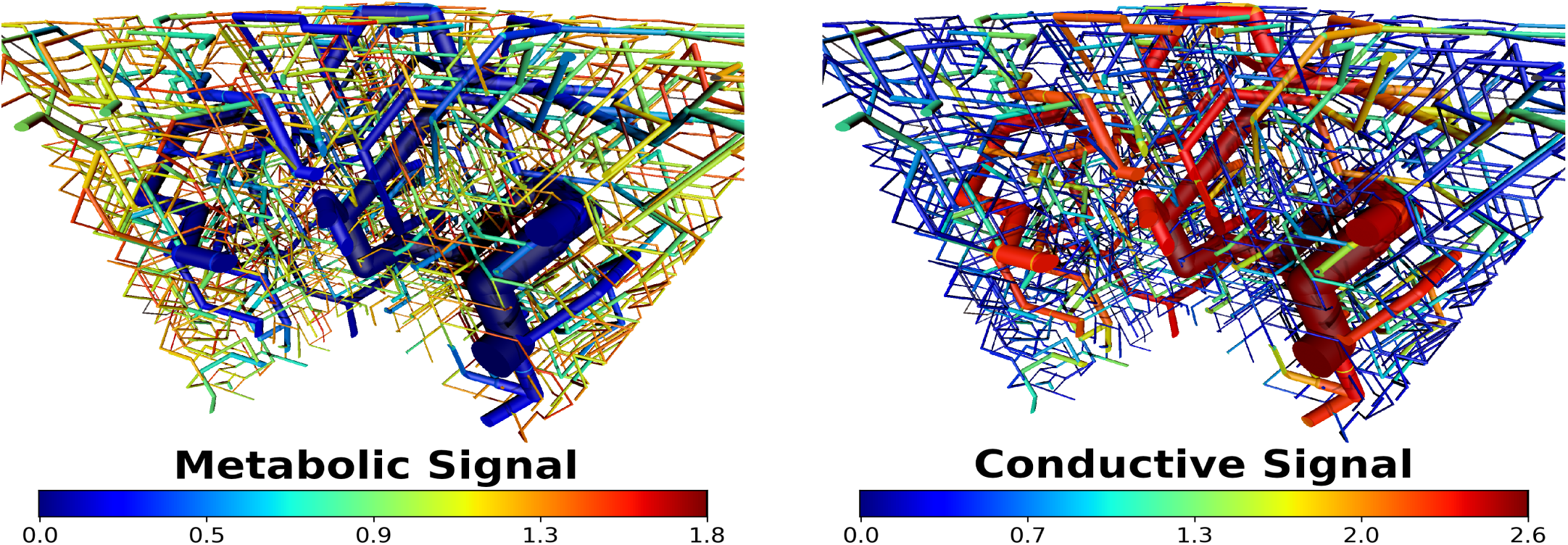
Adaptation signals (section 2.1) for one representative of the ensemble of synthetic arteriovenous networks. Left column: metabolic signal (S_3_) Right column: conductive signal (S_4_)

### 3.4 Murray’s law

The networks created by Tumorcode fulfill Murray’s law by construction (section 2.2). In the previous section, we applied the adaptation algorithm to them and investigated the resulting differences between the networks before and after vessel adaptation. Here we check for a single network (shown in figure 8) whether it fulfills Murray’s law.

We plot the cubic root of the sum of daughter vessel radii cube against the radius of the mother vessel for every intersection in figure 10. Remarkably, the data is oriented along the diagonal meaning the relations between mother and daughter vessels are close to Murray’s law. Given the radius of two daughter vessels, Murray’s law (equation 1) calculates the mother vessel radius. The recursive application of Murray’s law within a network of arbitrary number of intersections leads to a complicated dependency similar to the dependency of the signal S_4_ in the adaptation algorithm. The distribution of differences between two daughter vessels at an intersection reveals the differences between Murray’s law and the vessel adaptation (figure 11). For the adaptation algorithm, the distribution is smooth and lacking discrete peaks. Since the network construction algorithm of Tumorcode uses a fixed capillary radius and a discrete vessel length (on a lattice), repeated application of Murray’s law leads more often to the same proportions at an intersection.

**Figure 10:**
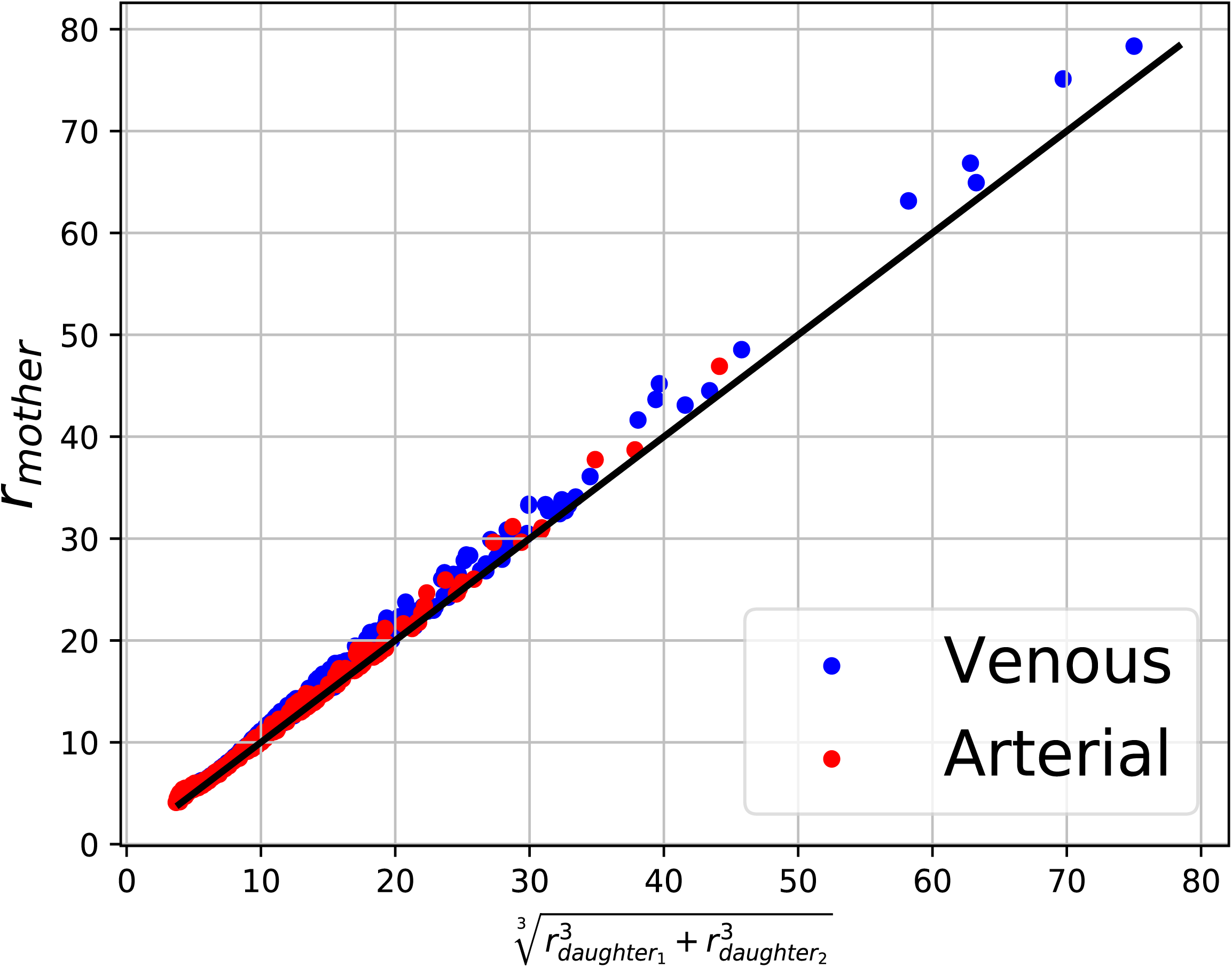
The black diagonal represents murray’s law for *α* = 3. The data points are taken from the A-network shown in figure 8 in the right column. Blue dots are venous intersections (431) and red dots are arterial intersections (475).

**Figure 11:**
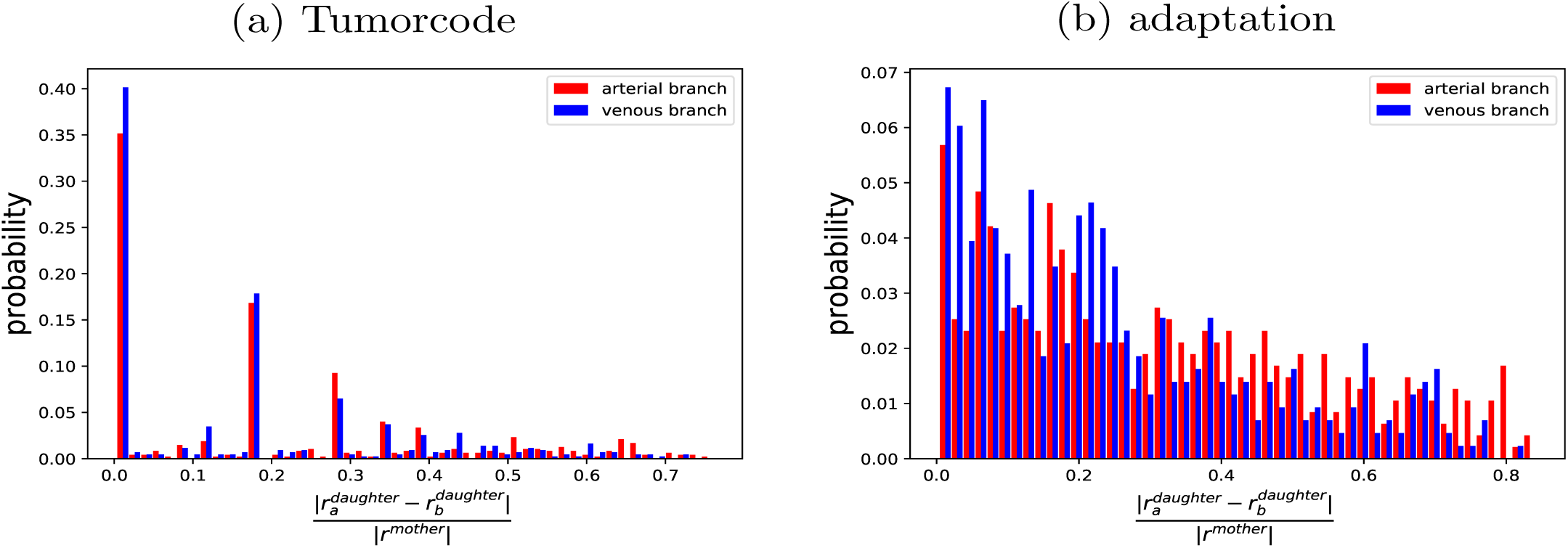
Normalized distribution of differences between two daughter vessels at a Y-shaped intersection. The used topology is the same for both cases and shown in figure 8. Left column: before adaptation (vessel radii fulfilling Murray’s law by construction in Tumorcode). Right column: same topology as left column but vessel radii adapted with parameter set listed in table 2.

We conclude that the adaptation algorithm together with our optimization objective (homogeneous flow inside the capillary bed) recovers Murray’s law.

### 3.5 Application to cancerous networks

The adaptation algorithm accounts for the modification of radii in blood vessel networks without topological modifications. In solid tumors, topological modifications are an essential part of the growth procedure and therefore considered in our simulation framework [21]. In [24], we simulated the oxygen distribution in breast carcinoma and found that an additional vessel dilation is required to obtain oxygen distributions in agreement with the clinical measurements. We conjectured that the adaptation algorithm could possibly provide a physiological mechanism for this vessel dilation. Indeed observe the expected behaviour in a minimal example (figure 12): the high metabolic demand (*HQ* < *Q*_ref_) of the vessels created by angiogenesis increases the radii during adaptation (see zoom in the upper right of figure 12). The new tumor vessels couple two previously uncoupled branches and thereby raise the topological signal in the connected branches as intended by the adaptation algorithm (see value of main inlet and outlet in bottom right of figure 12). Although this works in this simplified example, we found that such modifications destabilize the convergence of the adaptation algorithm when occuring in a repeated manner or at multiple locations at the same time as it happens our simulations of tumor growth in large arteriovenous systems.

**Figure 12:**
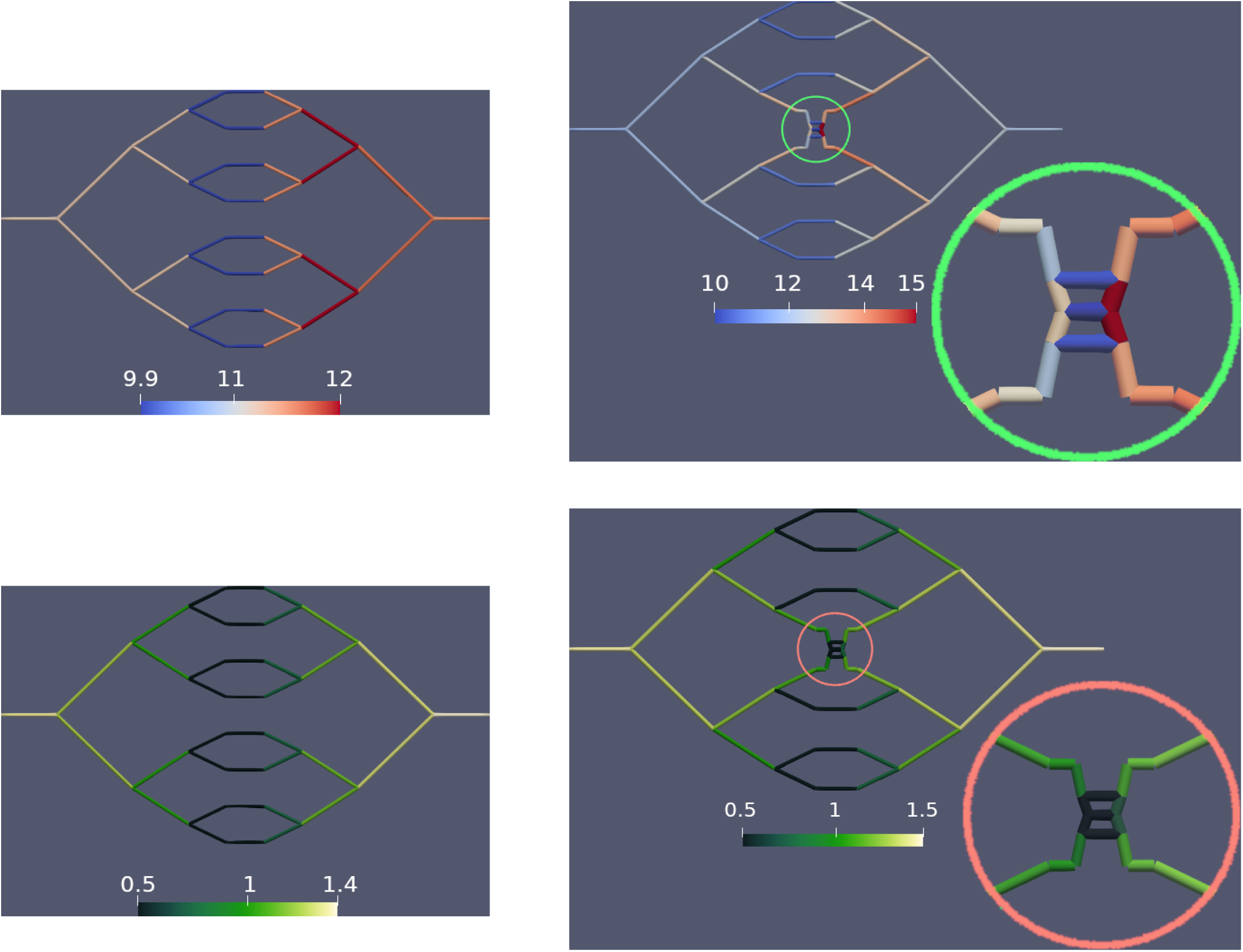
Critial behaviour during angiogenesis. We constructed a minimal example consisting of four closed capillary loops which is representative for the situation larger arteriovenous networks. Left column: w/o tumor, right column: slightly remodelled situation by angiogenesis during tumor growth with zoom into the critical area. Top row: vessels color-coded by their vessel radius. Bottom row: vessels color-coded by their value of the conductive signal. The complete example with the topology, the boundary conditions and the adaptation parameters is documented in the source code distributed with this manuscript or on GitHub at https://github.com/thierry3000/tumorcode (py/krebs/adaption/testAdaption.py)

## 4 Discussion

Pries et al. constructed an algorithm to dynamically regulate vessel radii within a blood vessel network. Whereas they used data from very rare animal models to fix the free parameters in their theory, we applied in this paper the adaptation scheme to artificial blood vessel networks and obtained the free model parameters without referring to experimental data (compare table 1). Instead, a computational optimization procedure (section 2.3) was used to estimate the parameters under reasonable assumptions which are a) biological boundary conditions (inflow and pressure at outlet) and b) homogeneous distribution of blood flow within the capillary bed.

We compared arteriovenous networks constructed by Tumorcode (**M-networks**, radii obey Murray’s law by construction) with networks having the same topology, but with vessel radii obtained by the adaptation algorithm (**A-networks**). From figure 8, figure 9, the data in section 3.2 and 3.3, it becomes apparent how the the two types differ. A quantitative analysis of regional blood volume (rBV, figure 5a and 6a) and surface to volume ratio (s2v, figure 5b and 6b) showed that the **A-networks** carry more blood while the **M-networks** offer a higher surface to volume ratio — especially at the capillary level.

The vessel radii of the **A-networks** are in agreement with Murray’s law which is derived by minimizing the energy required to overcome the viscous drag and to maintain a constant volume flow (section 3.4). Drag and flow are not directly used in the adaptation model, but its signals involve a form of electrical or topological energy (conductive signal), chemical energy (metabolic signal) and the tension of endothelial layer (shrinking tendency). In Tumorcode, we consider also the pathological case where a tumor affects the vasculature by wall degeneration, dilation, regression, collapse and angiogenesis. We tried to include the adaptation algorithm as additional element to alter the blood vessel radius in the Tumorcode software. Providing a convergent set of adaptation parameters for a given arteriovenous network (as found in this manuscript), the convergence was not achieved when tumor growth induced angiogenesis and vessel collapse are present. The discussion of the stability of networks with low generation shunts in the appendix of [17] illustrates why the algorithm cannot be stable without the conductive signal. But even including the conductive signal, the adaptation algorithm becomes unstable for a single set of parameters when the network topology is subsequently altered (section 3.5). It should be noted that some mathematical models of vascularized tumor growth include vessel radius adaptation [26], but no conductive signal, although it is an integral component of the vessel adaptation theory.

The signals introduced by the adaptation algorithm are based on biological relevant processes which provide an input for mathematical modelling. Currently the signals are modeled with parameters (compare table 1), but in future the drop in voltage along a vessel or its metabolic uptake might become experimental input fixing the free parameters of the adaptation model.

## Supporting information

Simulation Details

## Acknowledgment

We thank Adam Wysocki for useful discussions and the DFG (German Research Foundation) for financial support by SFB 1027.

## Data availability

We support open science by sharing our software and the created raw data at www.zenodo.org.

### Software

A developer version of the Tumorcode software tool [21] is hosted at www.github.com/thierry3000/ For the presented manuscript, we worked with the Pre-release v1.1.0-alpha.1 (Adaptation) version of the code which we also archived [27].

### Data

We store the results of executed simulations, the used settings and parameters, the network topologies and the images in a single compressed archive [28].

