## Supplementary material for "Dynamic vessel adaptation in synthetic arteriovenous networks": Simulation Details

### Details of the particle swarm optimization as reported in "Dynamic vessel adaption in synthetic arteriovenous networks"

Thierry Fredrich

2019

Our aim was to use deap (Distributed Evolutionary Algorithms in Python) to spread the evaluation of multiple vessel adaptations with different boundary conditions and algorithm parameters across multiple processors.

#### 1 Required Software

We used the Intel Parallel Studio XE 2018 with its compiler to obtain fast executable. This fact required us to make us of the IntelPython suite. We installed *libzmq* and the python bindings from source.

This enables us to install SCOOP (Scalable COncurrent Operations in Python) that is nice module to distribute computational task from heterogeneous grids to supercomputers. In our case we used a small (64 Nodes with up to 28 cores) beowulf cluster hosted by the theoretical physics department of Saarland University and payed by SFB 1027. Apart from the aforementioned software the cluster runs operationg system Gentoo linux hardened SLURM queuing system.

#### 2 Particle swarm optimization

We conducted Particle swarm optimization with the deap software based on their example in the documentation. Our source code is available on GitHub at: https://github.com/thierry3000/tumorcode. (py/krebs/adaption).

All optimizations listed in table 1 were conducted with a population of 320 individuals and 10 generations of mutation. We used 20 Nodes with 16 cores each to have 320 independent tasks. For the discussion presented in the main article, we evaluated the PSO with the parameter set value_list_14 however we performed more optimizations. All successful runs could be downloaded at: .

| name | #boundary conditions | flow/$10^7$ $\mu m^3/s$ | pressure/ kPa |
| --- | --- | --- | --- |
| value_list_11 | 160 | 1-9 | 3.5-4.5 |
| value_list_12 | 160 | 1-9 | 3.5-4.5 |
| value_list_13 | 864 | 1- 90 | 1.8-4.5 |
| value_list_14 | 342 | 0.5-10 | 0.5-9.5 |

Table 1: Datasets resulting from successful PSO (downloadable at: ). All optimizations were conducted with a population of 320 individuals and 10 generations of mutation.
